# Natural transformation of the filamentous cyanobacterium *Phormidium lacuna*

**DOI:** 10.1101/870006

**Authors:** Fabian Nies, Marion Mielke, Janko Pochert, Tilman Lamparter

## Abstract

Research for biotechnological applications of cyanobacteria focuses on synthetic pathways and bioreactor design, while little effort is devoted to introduce new, promising organisms in the field. Applications are most often based on recombinant work, and the establishment of transformation can be a risky, time-consuming procedure. In this work we demonstrate the natural transformation of the filamentous cyanobacterium *Phormidium lacuna* and insertion of a selection marker into the genome by homologous integration. This is the first example for natural transformation of a member of the order Oscillatoriales. We found that *Phormidium lacuna* is polyploid, each cell has about 20-100 chromosomes. Transformed filaments were resistant against up to 15 mg/ml of kanamycin, and the high resistance feature allowed for rapid segregation into all chromosomes. Formerly, natural transformation in cyanobacteria has been considered a rare and exclusive feature of a few unicellular species. Our finding suggests that natural competence is more distributed among cyanobacteria than previously thought. This is supported by bioinformatic analyses which show that all protein factors for natural transformation are present in the majority of the analyzed cyanobacteria.

## Introduction

Biotechnology oriented research with cyanobacteria ranges from the production of low-cost material like bulk chemicals or biofuels (1) to high-value compounds like pharmaceutics (2). Advancements in cyanobacterial biotechnology are based on continued optimization of photobioreactors, the introduction and improvement of metabolic pathways by recombinant DNA technology, and the search for suitable organisms (3, 4). The establishment of protocols for gene transfer can be challenging for new cyanobacteria, because of barriers like extracellular materials and nucleases (reviewed in (5)). There are three common methods for gene transfer into cyanobacteria: electroporation, conjugation, and natural transformation (NT). For NT, cells have to be in a physiological state, termed natural competence, in which the recipient cell is able to actively transport DNA into the cytoplasm. Protocols for NT are generally simple and straight forward (6, 7), but only few naturally competent cyanobacteria (NCC) are known: diverse *Synechococcus* (8, 9) and *Synechocystis* (10, 11) strains, *Microcystis aeruginosa* PCC 7806 (12) and *Thermosynechococcus elongatus* BP-1 (13). These cyanobacteria have a unicellular lifestyle, and NT was frequently described as a unique feature of few unicellular cyanobacteria (6, 14, 15). We also found one report about NT of the filamentous cyanobacterium *Nostoc muscorum* (16), but are not aware of other filamentous cyanobacteria.

Natural transformation in bacteria is dependent on pil proteins of type IV pili. Furthermore, the competence proteins ComEA, ComEC, and ComF as well as the DNA processing protein DprA and the DNA recombination and repair protein RecA are essential (17-19). For the cyanobacterium *Synechocystis* sp. PCC 6803, *comEA, comF, pilA1, pilB1, pilD, pilM, pilN, pilO, pilQ*, and *pilT1* knockout mutants are deficient in NT (20-23). Although in a recent survey, homologs of these and other competence related genes were found in most cyanobacterial genomes (24) there is so far no experimental evidence for NT in a novel species since more than a decade.

Most cyanobacteria possess multiple chromosomes per cell (25-29). After transformation, homologous recombination results in the integration into one chromosome. Segregation into the other chromosomes can be achieved by selection on increasing antibiotic concentrations. Until complete segregation is achieved, the transformants may be unstable and the integrated sequences can be lost again under nonselective conditions (30, 31).

In this work we established an NT protocol for *Phormidium lacuna*, a filamentous cyanobacterium of the order Oscillatoriales, a species isolated and characterized by our workgroup as a promising candidate for biotechnological applications (32). This is to our knowledge the first transformation protocol for the genus *Phormidium* and the first report of NT for the order Oscillatoriales. *Phormidium lacuna* was transformed by the integration of the kanamycin (Kn) resistance cassette (*kanR*) into the genome via homologous recombination. Clones were selected by Kn resistance and integration into genome was validated by PCR. During clone validation it was found that *Phormidium lacuna* is polyploid. This was confirmed by a DAPI fluorescence assay. By comprehensive BLAST analysis based on sequences of essential proteins for natural transformation (natural transformation factors, NTFs) we predict that a large fraction of cyanobacteria might be naturally transformable.

## Methods

### Plasmids for transformation

For transformation studies, we constructed plasmids with a *Phormidium lacuna* sequence sc_7_37 (position 37 in DNA scaffold 7) that is interrupted in the middle by a kanamycin resistance cassette. A 1138 bp and a 2167 bp product were generated by PCR using Q5 polymerase (NEB, Ipswich, MA, USA) and HE10DO DNA as template. The primer pairs were t256: CGTGCGAGACTCAACCCAAAC / t257: GAAACCTGATCGAACCGTTTTAC for the short sequence and F114: TTGTTCGAGGCAGTTGCG / F115: TGACAATGGGGTGGAGGG for the long sequence. The sequences were integrated into pGEM-T (Madison, WI, Promega, USA). The *Phormidium lacuna* sc_7_37 sequence encodes for a putative hydrogenase (WP_087706519); the sequence of the strain used in the present study, HE10DO, is identical with *Phormidium lacuna* HE10JO for which the genome sequence is established (32). pGEM-T plasmids are not propagated in cyanobacteria due to incompatible origin of replication (33).

The kanamycin resistance cassette kanR was PCR amplified from pUC4K (34) using the primer pair GG1: CAACAAGAAGACGGAACCTAGGCACCCCAGGCTTTACAC / GG2: CAACAAGAAGACGCAAACTTTGCTTTGCCACGGAACGG. The plasmid with the long sc_7_37 insert was amplified with the primers GG3: CAACAAGAAGACCCGTTTGCGAGGCTAAAGGC / GG: 4 CAACAAGAAGACACGGTTCCCACTCCCAAAGC. The resulting plasmid pFN_7_37_2k_kanRn was generated by the type IIS restriction enzyme BbsI and T4 DNA ligase (both NEB, USA). Another version of kanR with slightly different 5’ and 3’UTR was generated by PCR using the primer pair F13: CAACAATCTAGACTCGTATGTTGTGTGGAATTG / F14: CAACAAGCTAGCCAAGTCAGCGTAATGCTCTG. This cassette was inserted in the plasmids with the long and the short homologous sc_7_37 sequence. Plasmids were amplified with the primer pair F5: CAACAAGCTAGCGTTTGCGAGGCTAAAGGCG / F6: CAACAATCTAGAGGTTCCCACTCCCAAAGC and DNA sequences digested with XbaI and BmtI (NEB, Ipswitch, MA, USA) and ligated. The resulting plasmids are termed pFN_7_37_kanR and pFN_7_37_2k_kanR plasmids for transformation were purified using Macherey Nagel (Düren, Germany) midi-prep plasmid purification kit.

### Cultivation of *Phormidium lacuna*

*Phormidium lacuna* strains HE10DO and HE10JO were cultivated at 23 °C in f/2 salt water medium (35, 36) or in f/2+ (in which nitrate and phosphate are 10x increased) under permanent illumination m^-2^ s^-1^ (30 µmol white light from fluorescent tubes Lumilux-DeLuxe L 18/954, Osram, Munich, Germany) and continuous shaking (70 rpm). For agar plates, f/2 medium without Na_2_SiO_3_ and with 1.5% Bacto Agar (BD Diagnostics, Franklin Lakes, NJ, USA) were used.

### Transformation

For transformation, *Phormidium lacuna* HE10DO (32) were cultivated in 100 ml f/2 medium to an optical density OD_750 nm_ of 0.25 - 0.35. The cell suspension was homogenized using an Ultraturrax (Silent Crusher M. Heidolph, Schwabach, Germany) with the dispersion tool 18F at 10,000 rpm for 3 min. The cell suspension was centrifuged at 6000 g and 4°C for 15 min. After each centrifugation step, the supernatant was removed. Cells were resuspended in 20 ml water (4 °C) and centrifuged again. This washing step was repeated. The cells were finally suspended in the residual liquid, transferred into 1.5 ml tubes and centrifuged again at 6000 g at 4°C for 15 min. Cells were finally suspended in 1 ml supernatant. Portions of 100 µl were mixed with 3-30 µg DNA in 10 µl water, transferred into 10 ml f/2 medium and cultivated for 2 d. Cells were again centrifuged, resuspended in 1 ml medium and transferred to f/2 agar plates with 0, 70 and 120 µg/ml Kn. Resistant lines were selected after 10 -28 d. Transgenic cells were cultivated on increasing Kn concentrations in f/2+ suspension culture until complete segregation was achieved. Electroporation experiments were performed with both strains HE10DO and HE10JO, whereas natural transformation was performed with the strain HE10DO only.

### Validation of transformants

To test for homologous integration, ca. 10 mg cell samples were homogenized and lysed mechanically by micropestle and subjected to PCR with Taq Polymerase (NEB, USA). The following primers were used: F25: GGTCTAGGTGAGGCAATCC / F28: ACCTGATTTGTTTATATCTGACGC for pFN_7_37_kanR transformants and F120: GGGTAGCCTAGACTCATCC / F121: ATGCGGAAGTGACTGAGG for pFN_7_37_2k_kanR and pFN_7_37_2k_kanRn transformants. All primers bind only in the *Phormidium lacuna* genome, upstream or downstream of the integration site.

### Bioinformatics

Protein sequences of the NTFs of *Synechocystis* sp. PCC 6803 were obtained from the NCBI data base. NTF sequences of 6 naturally competent cyanobacteria (NCC) were identified by the offline NCBI tool BLAST+ (version: 2.7.1, (37)) based on the annotated NTFs of *Synechocystis* sp. PCC 6803. All NTFs identified in this way were used in a BLAST query against all cyanobacterial sequences of the NCBI non-redundant database (August 2018). Bit score as indicator of homology was processed by a minimum homology quotient method: For each homolog, the bit score of each alignment was divided the by smallest bit score of the respective NCC pairs. This value is given in Table 3.

**Table 1.**
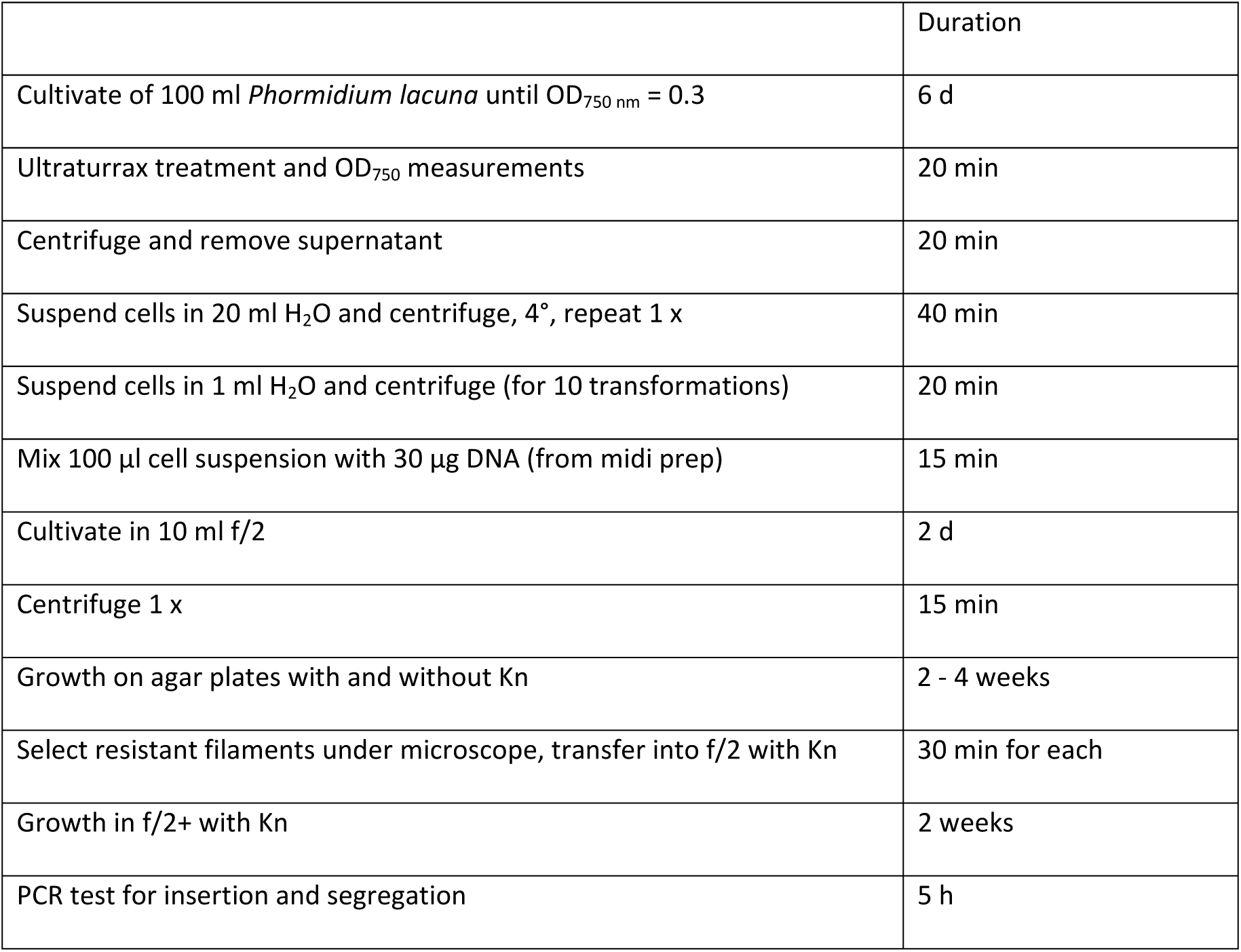
Summary of NT protocol for *Phormidium lacuna*. Additional details are given in the methods section.

**Table 2.**
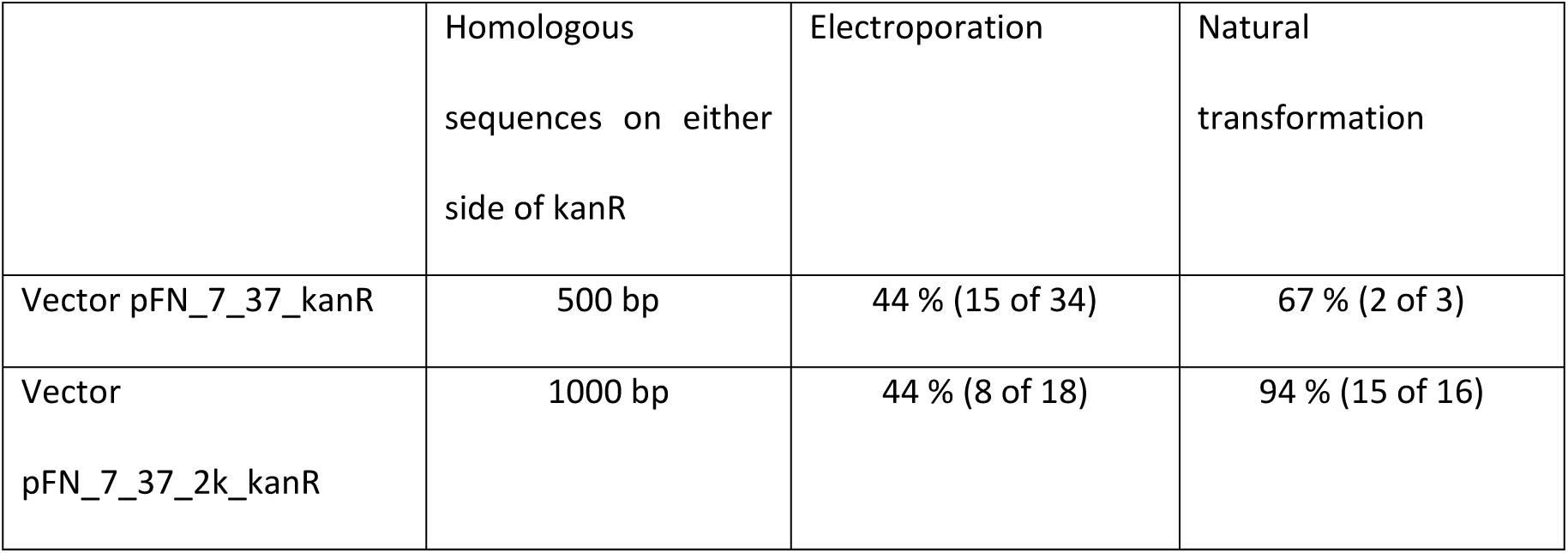
Transformation of *Phormidium lacuna* HE10DO by electroporation and natural transformation. The numbers stand for successful transformations i.e. 1 or more resistant lines could be isolated in the relevant trial. Electroporation was performed in 1 mm cuvettes and 300 V pulses. Mixed plasmid DNA and cell suspensions were suspended in 1 ml medium and transferred into culture flasks directly after the electric pulse.

**Table 3.**
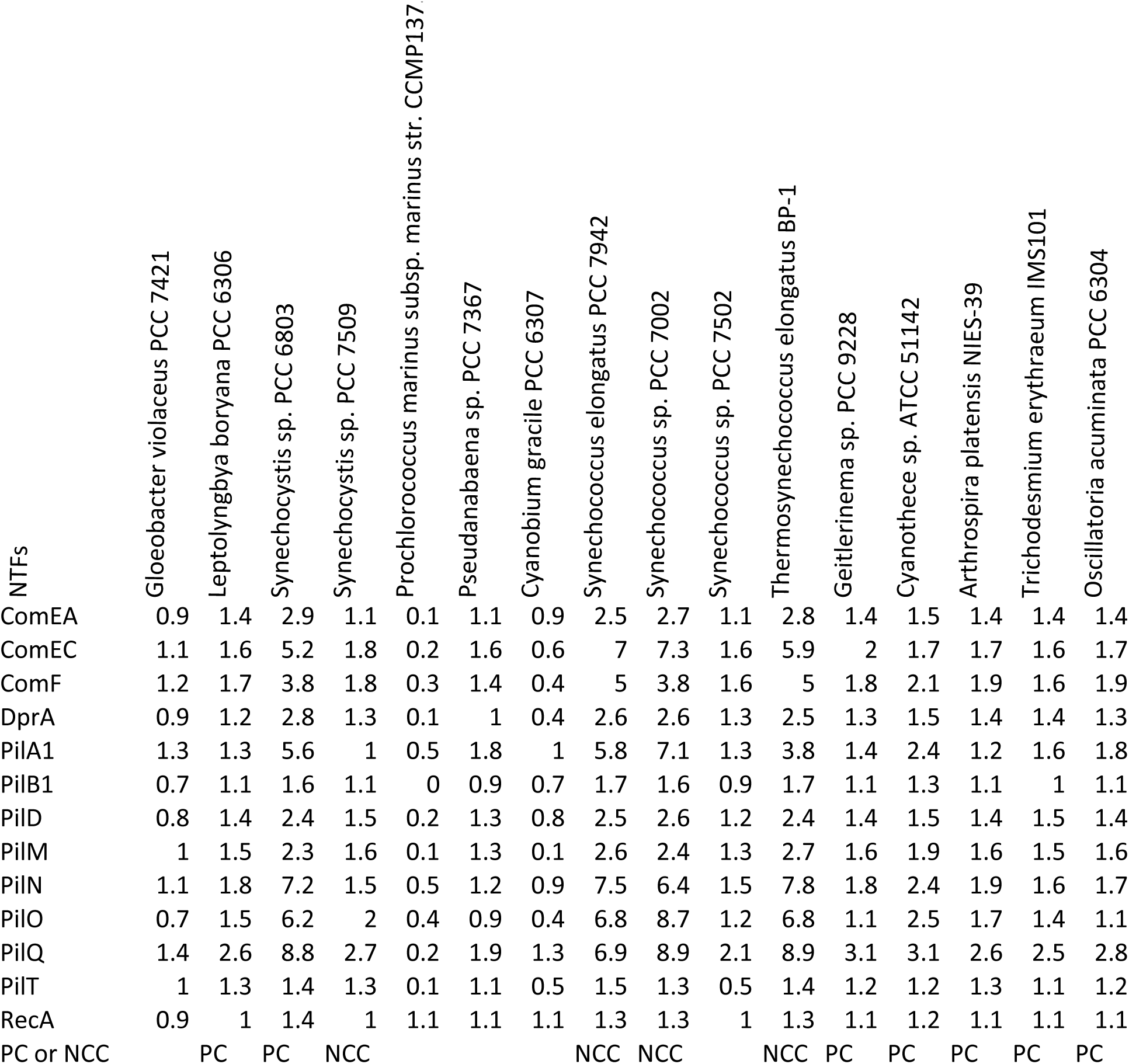
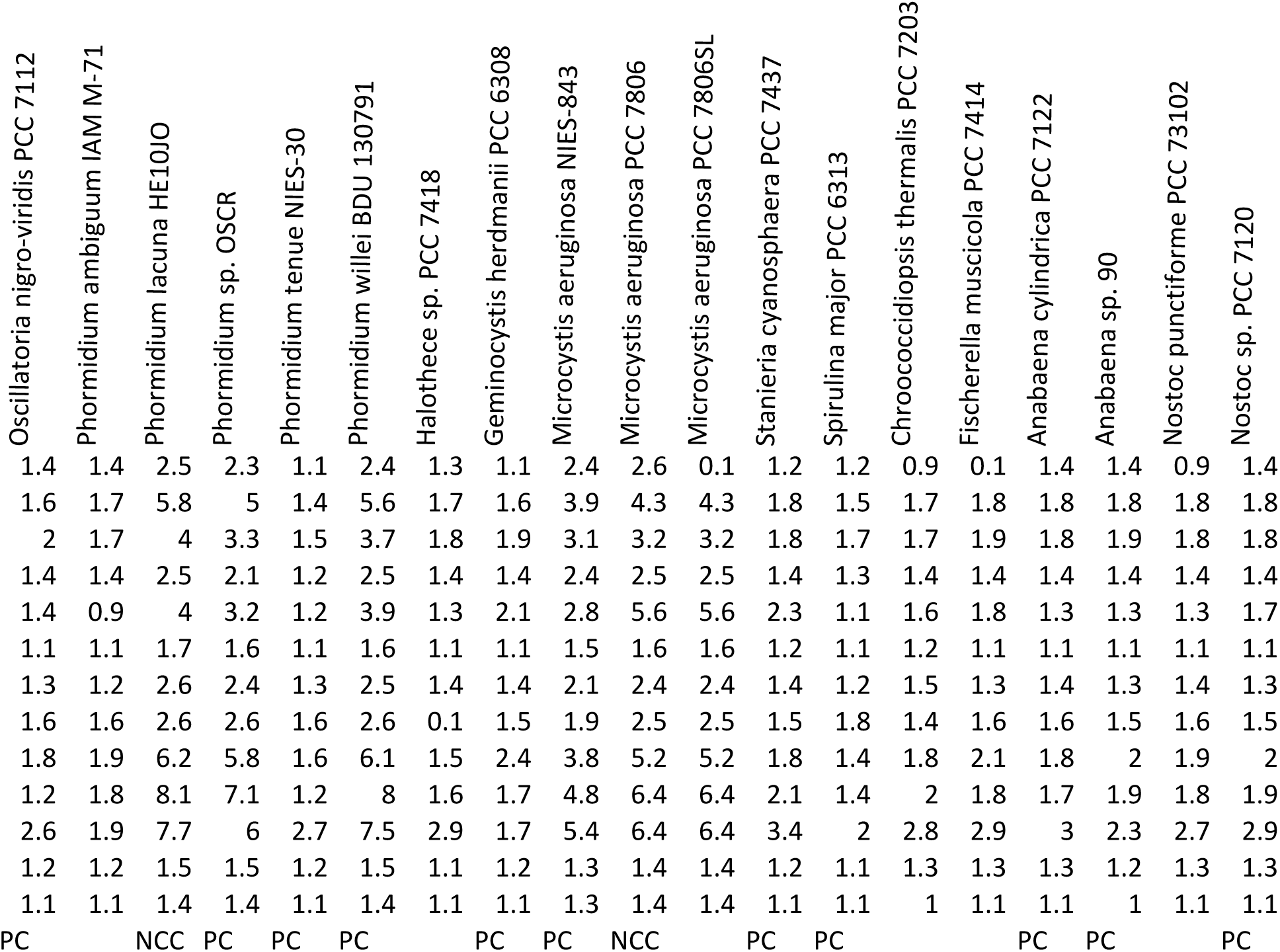
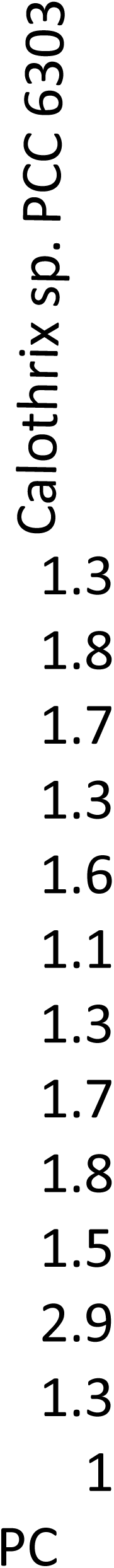
Summary of minimum bit score values of (putative) NTFs of 36 cyanobacterial species. In the first step, NTF homologs of 6 naturally competent cyanobacteria (indicated by NCC in the last line) were identified. These NTFs were used as queries in a BLAST search against protein sequences of 30 cyanobacterial species. The bit score of each homology hit was divided the by smallest bit score of the respective NCC pairs.

### Chromosome copy number

*Phormidium lacuna* liquid cultures were homogenized by an Ultraturrax for 3 min at 10,000 rpm. To determine the cell concentration, the total length of all filaments in a given volume of a counting chamber was estimated (about 50 filaments for each sample) and divided by the average cell length (4 µm). Cells were quantitatively disrupted using an Aminco French pressure cell (Thermo Fischer, Waltham, MA, USA). A fluorescence excitation spectrum from 300 to 400 nm was recorded at 490 nm emission. Thereafter, DAPI was added to a final concentration of 100 ng/ml. The fluorescence spectrum was measured again. After addition of 5 µl DNase (75 Kunitz) the fluorescence decreased over 2 h. The difference of peak values before and after DNA digestion was taken as measure for DNA concentration. As a reference, calf thymus DNA at various concentrations was dissolved in the same medium and the exact concentrations determined by A_260 nm_. The same fluorescence recordings were performed as with *Phormidium lacuna* DNA. Final values were corrected by GC contents of both species, since GC does not induce DAPI fluorescence (38). A genome size of 3.5 Mio bp from genome sequencing was taken for *Phormidium lacuna* (32).

## Results

We initially established a transformation protocol for *Phormidium lacuna* by electroporation that was based on protocols for related filamentous cyanobacteria (39-42). For integration into cyanobacterial DNA we used homologous recombination, which works at high efficiency in cyanobacteria (43) and other bacteria. In the transformation vectors, the homologous sequence “sc_7_37” was interrupted by a kanR resistance cassette. With these vectors, we obtained Kn resistant lines in about 40% of trials. During electroporation studies, we isolated a resistant line from a control experiment in which cells were incubated with DNA, but no electroporation pulse was given. The transformation protocol could be optimized (Table 2 and methods) so that in almost all transformation assays, DNA was integrated in *Phormidium lacuna* cells. Fig. 1 shows examples for filaments on selection medium at different time points after transformation.

**Fig. 1.**
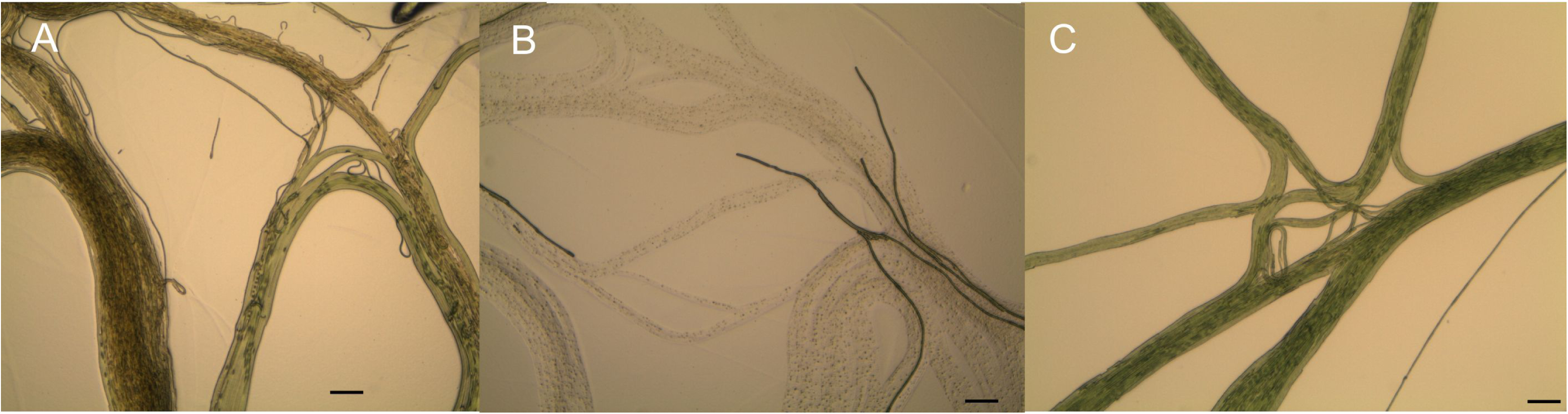
Filaments of *Phormidium lacuna* after transformation with pFN_7_37_kanR during or after selection on agar plates. A) 2 weeks after transformation and growth on 70 µg / ml Kn. Brownish and greenish filaments can be distinguished which represent non-transformed or transformed cells, respectively. B) 2 weeks after transformation and growth on 120 µg / ml Kn after, in this area of agar plate, living and dead filaments are clearly distinguished C) 4 weeks after growth on 120 µg/ml Kn, all filaments are resistant. Bundles of parallel filaments are characteristic for growth on agar, the growth pattern is comparable to that of wild type on Kn-free agar. Scale bars 100 µm.

During our studies, we used plasmids pFN_7_37_kanR and pFN_7_37_2k_kanR(n) (Fig. 2 A and methods) that differ by the length of the homologous sequence; the kanR resistance cassette is flanked by ca. 500 bp or 1000 bp homologous sequences on each side, respectively. In a comparison of electroporation and NT that were performed under similar conditions (Table 1 and methods section), the success rate was always higher for NT as compared to electroporation (Table 2). When the pFN_7_37_2k_kanR(n) vectors was used, 15 out of 16 NT transformation trials were successful, i.e. resulted in the isolation of resistant lines that could be confirmed by PCR (see below). In the electroporation experiments, only 44 % transformation trials were successful. We assume that the deleterious effect of the electric pulse overrides the positive effect that the pulse might have on DNA incorporation into the cells. For *Synechocystis* sp. PCC 6803 it was observed that longer flanking sequences are beneficial for the homologous recombination into the genome (44). For *Phormidium lacuna*, we found no significant difference of transformation success between 500 bp and 1000 bp flanking sequences (Table 2). The length of the insert can also be critical for integration into the genome. In further transformation experiments we used inserts up to a length of 4241 bp. These could also be integrated into the genome of *Phormidium lacuna*, although with lower transformation efficiencies. It should be noted that the electroporation experiments were performed with both strains HE10JO and HE10DO, whereas natural transformation experiments were only performed with HE10DO.

**Fig. 2.**
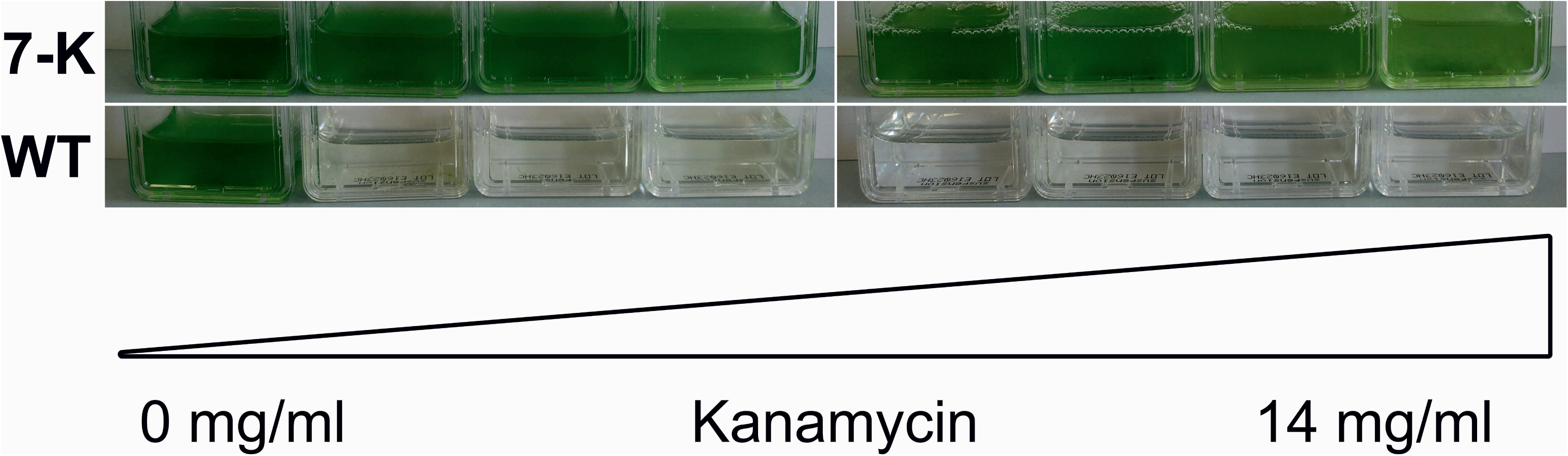
Kanamycin resistance of *Phormidium lacuna* HE10JO wild type and pFN_7_37_kanR transformants, liquid cultures 1 week after inoculation and growth under white light. Wild type cannot proliferate at 50 µg/ml Kn or above, while pFN_7_37_kanR transformants can grow up to 14.3 mg/ml Kn. Kanamycin concentration from left to right: 0, 50, 100, 200 μg/ml; 4.5, 8.3, 11.5, 14.3 mg/ml.

We found that transformants with the kanR cassette were resistant against very high Kn concentrations. Transformed lines could be routinely cultivated up to at least 15 mg/ml Kn (Fig. 2). In 50 mg/ml Kn, transformants remained green for few days but did not survive the first week. The upper limit for Kn resistance is thus around 20-40 mg/ml. *Escherichia coli* DH5α and *Synechocystis* sp. PCC 6803 cells, transformed with the same kanR, were resistant up to ca. 200 µg/ml and ca. 500 µg/ml Kn, respectively. In an extensive literature survey we found a report about an environmental *Enterococcus* strain that was resistant up to 2 mg/ml Kn (45), but no report about higher Kn resistance. Thus, transformed *Phormidium lacuna* has probably the strongest Kn resistance reported so far.

We wanted to integrate the resistance cassette into a neutral site in order to allow expression of additional introduced genes in future experiments. The selected homologous sequence codes for a hydrogenase homolog (WP_087706519). Cyanobacterial hydrogenases are oxygen sensitive (46) and should not be active under our standard growth conditions with continuous illumination. Indeed, the transformants did not display an observable phenotype.

The integration into the genome was validated by PCR. Figure 3 shows results from a transformation with pFN_7_37_KanR. The used primers bind to regions in the genome of *Phormidium lacuna* just upstream or downstream of the insertion site, respectively. In wild type extracts, an expected short PCR product of 1200 bp was detected (Fig. 3). Lines T1a and T1b are from filaments that had been cultivated in liquid medium with 1000 µg / ml kanamycin for 4 weeks after transformation and the expected long PCR product with 2600 bp was detected. In T2a and T2b, which are from filaments that were cultivated for 9 d in liquid culture with 250 µg/ml Kn, both the 1200 bp and the 2600 bp PCR products were present. Such a “partial integration” is known from transformations of other cyanobacteria where it results from the polyploid character of these cells (25). In our *Phormidium lacuna* transformation experiments, the pattern of partial integration during early selection and complete integration after prolonged selection was observed in 8 independent experiments. In order to find out whether *Phormidium lacuna* cells are also polyploid, we estimated the number of chromosome copies by a DAPI based fluorescence assay. In single experiments, the estimated values of chromosome copies varied between ca. 20 and ca. 120. *Phormidium lacuna* has thus multiple chromosome copies per cell. We assume that the large variations are not only due to methodological variations, since in control measurements with calf thymus DNA the variations were much smaller. Although cultivation was performed under standardized conditions, yet unknown factors must affect the chromosome copy number of whole samples. However, when the *Phormidium lacuna* data were plotted against cell densities (Fig. 3), values at lower densities i.e. shorter time of propagation, correlated with higher numbers of chromosome copies; at higher OD, the copy numbers were around 20. Thus, there is an influence of growth phase on the chromosome copy number in *Phormidium lacuna*, which has also been reported for *Synechocystis* sp. PCC 6803 (24, 28).

**Fig. 3.**
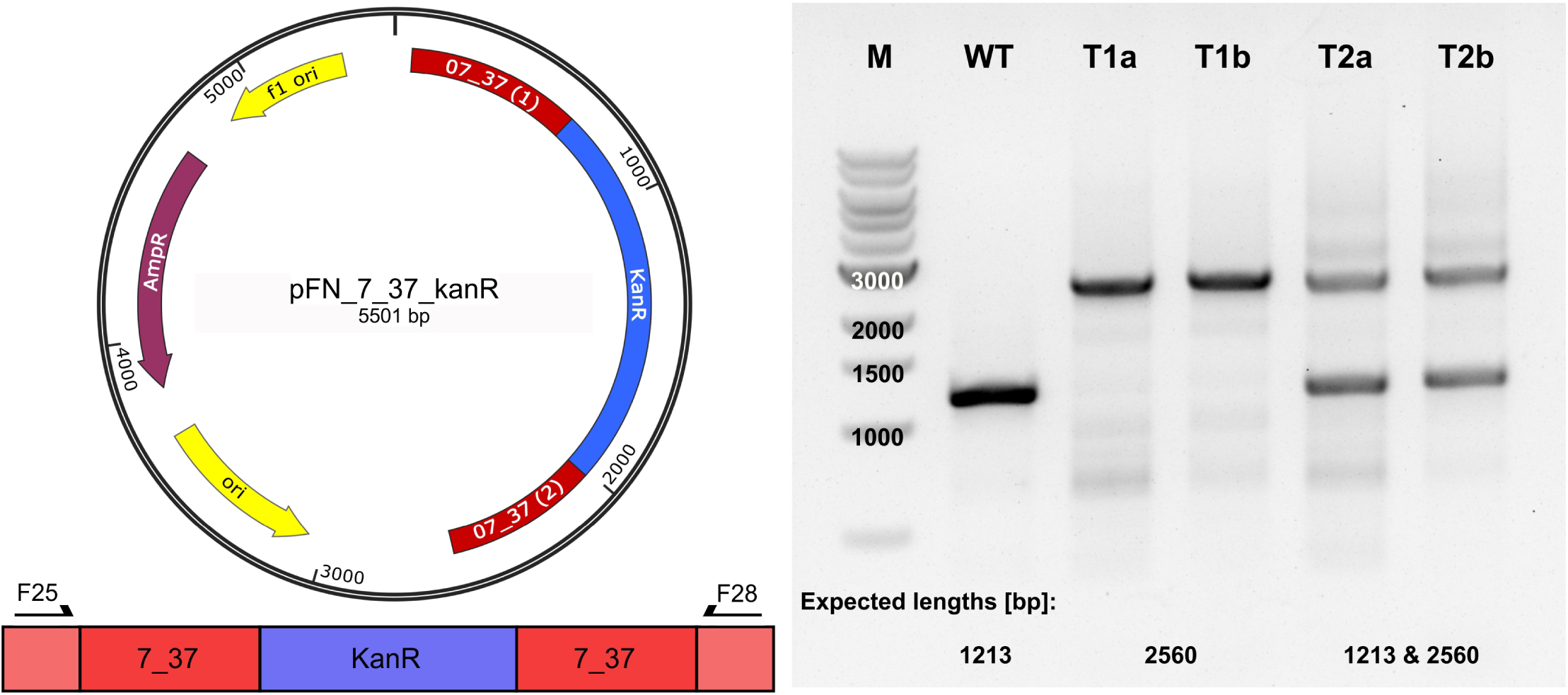
Validation of *Phormidium lacuna* transformants by PCR. a) Plasmid map of pFN_7_37_kanR, red - homologous sequences of the sc_7_37 locus, blue - kanamycin resistance cassette kanR, yellow - origin of replication (f1 - bacteriophage origin, other - pUC origin for *E. coli*), purple - ampicillin resistance cassette. b) Integration site of pFN_7_37_kanR and primer binding sites. red - homologous sequences encoded on the vector, pale red - *Phormidium lacuna* chromosome, blue - kanamycin resistance gene. Primer pair: F25/F28 covering whole insertion site. c) Agarose gel for the PCR with the primer pair that covers the full insert for *Phormidium lacuna* WT and transformants. Transformants T1a and T1b were cultivated after selection on agar plate for one cultivation period (7 days) in f/2+ liquid medium with 250 µg/ml Kn and one period in f/2+ with 1000 µg/ml Kn. T2a and T2b were cultivated only for one period at 250 µg/ml Kn. M: 1kb DNA ladder (NEB, USA). Integration of the kanR cassette into the genome of *Phormidium lacuna* is indicated by the larger PCR product (2560 bp).

We also studied by PCR how the Kn concentration in the medium affects the segregation of kanR in the genome (Fig. 4). After transformation and selection of a resistant line and one cultivation cycle in suspension culture with 100 µg/ml Kn, the filaments were divided and cultured at 0, 100, 980 and 8300 µg/ml in suspension culture with subcultivation every 7 d for 4 weeks. After the first subcultivation, two PCR bands were observed in all cultures, indicating that Kn resistance was integrated in a part of the chromosomes but not in all (Fig. 4 a). Without antibiotic pressure, the 2600 bp PCR product decreased transiently and increased again after the 4^th^ subcultivation, and the 1200 bp wild type PCR product was present through all subcultivations. In the 100 µg/ml Kn samples, the wild type band was diminished after the third subcultivation and almost, but not completely, lost after the 4^th^ subcultivation. The results were similar for the selection on 980 µg/ml Kn. With 8300 µg/ml Kn, the wild type band disappeared almost completely already after the 2^nd^ subcultivation and was apparently lost after the 3^rd^ and 4^th^ subcultivations. Thus, high concentrations of Kn result in rapid segregation of kanR into all chromosomes. The high Kn resistance of *Phormidium lacuna* transformants could therefore provide an advantage for fast segregation of the selection marker in comparison to other cyanobacteria (47).

**Fig. 4.**
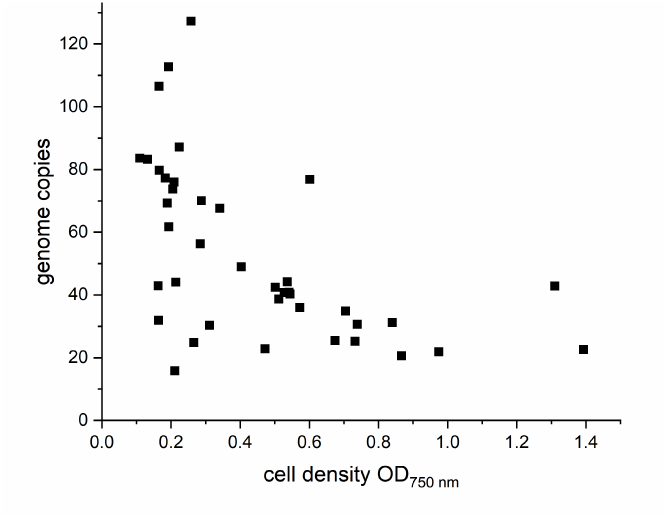
Chromosome copies of *Phormidium lacuna* cells estimated by DAPI fluorescence of filaments grown 1-5 days after inoculation. Each data point represents one independent measurement as described in detail in the methods section. Values are plotted against OD_750 nm_.

**Fig. 5.**
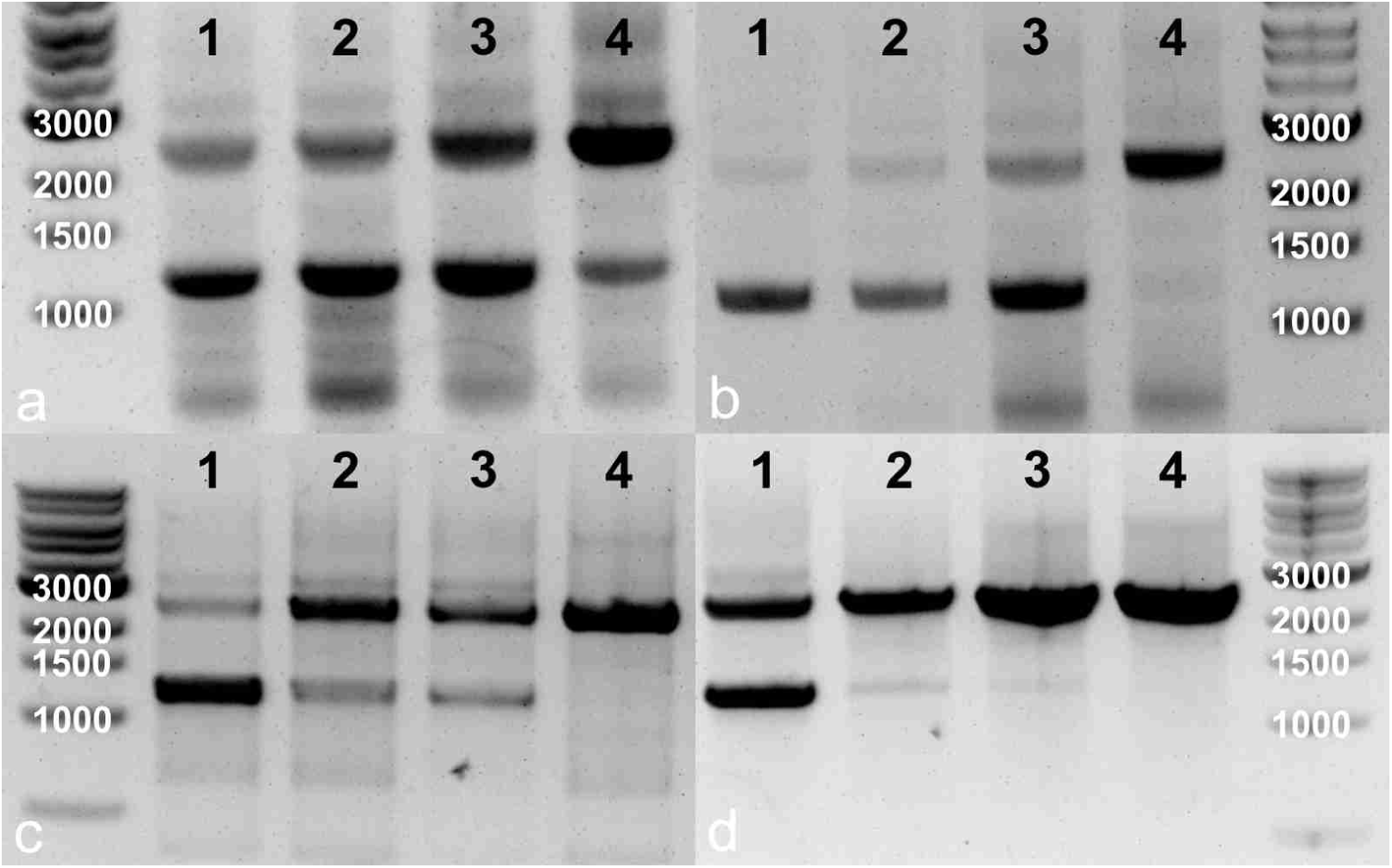
Detection of wild type and recombinant chromosomes in Kn resistant *Phormidium lacuna* pFN_7_37_kanR lines. Following cultivation in 100 µg/ml Kn until 2 weeks after transformation, the sample was divided and subcultivated one to four times on the Kn concentrations given in the panel. Primers: F25, F28. Marker: 1 kb DNA ladder (NEB, USA).

*Phormidium lacuna* HE10DO is the first representative of the order Oscillatoriales for which a NT protocol was established. According to mutant studies with *Synechocystis* sp. PCC 6803, proteins of the type IV pili **PilA1, PilB1, PilD, PilM, PilN, PilO, PilQ, PilT1** and the DNA receptor **ComEA** and **ComF** are NTFs, i.e. required for NT (20-23). Other NTFs that are essential for NT in general (but loss of NT has not been demonstrated in cyanobacteria) are ComEC, DprA and RecA (19). A functional type IV pilus in combination with this set of expressed non-pili proteins is probably not only crucial but also sufficient for NT. We can therefore assume that the probability for NT is high if a species has functional homologs of all proteins. In order to get an overview about the distribution of these proteins in selected cyanobacteria, we first identified NTF BLAST homologs in the naturally competent cyanobacteria (NCC), *Synechocystis* sp. PCC 6803, *Synechococcus elongatus* PCC 7942, *Synechococcus* sp. PCC 7002, *Thermosynechococcus elongatus* BP-1, *Microcystis aeruginosa* PCC 7806 and, *Phormidium lacuna* HE10JO (8-13). The strains *Phormidium lacuna* HE10JO and HE10DO are very similar and the genomic data for HE10JO is considered also relevant for the strain HE10DO (31). This set of proteins is used to define a range of similarity to decide whether other homologs are functional or not. We BLASTed all NTFs of all NCCs against the genomes of selected other cyanobacteria. In order to normalize each target, the highest bit-score (out of 6) was divided by the lowest bit-score among the 6 NCCs (i.e. the most unrelated NTF pair among NCCs). If the quotient is >= 1, we regard the target protein as functional homolog, because it is then within the range of NCC. If all NTF homologs of a species have values >= 1, the chances are high that NT is possible. Among 30 cyanobacterial species for which no NT is reported, 19 have quotients >=1 for all 13 NTF homologs and are thus promising candidates for NT. Among them are 14 filamentous cyanobacteria including two members of the genera *Arthrospira / Spirulina* and *Trichodesmium* with high economic impact. For other species the quotient for one or more NTFs is below 1, yet all essential genes for NT seem to be present in the genome but in lower homology to the NCC. These organisms may also be naturally transformable but it may be less likely due to the homology based prediction. Quotients ≤ 0.2 are considered as indicative for random hits during the BLAST search. Therefore, the 5 species that have at least one quotient ≤ 0.2 are regarded as critical for NT.

## Discussion

Even though natural competence is known for several single celled cyanobacterial species and one filamentous cyanobacterium, *Nostoc muscorum* (16), this mechanism was considered as rare trait among cyanobacteria (6, 14, 15). The finding of natural competence in *Phormidium lacuna*, the first example for the order Oscillatoriales, was therefore unexpected and surprising. Since now a new order of cyanobacteria is addressed, we assume that natural competence is broadly distributed among cyanobacteria. Our bioinformatic studies on the distribution of NTF homologs among 30 species, which are intended to be a representative selection of the cyanobacterial phylum, supports this hypothesis: functional homologs of all NTFs are present in at least 19 species. Thus, NT seems to be a widely distributed physiological function in cyanobacteria and could contribute more to evolutionary adaptations than previously suggested. It might play a major role in horizontal gene transfer, genetic recombination, and DNA repair in this phylum. Natural transformation is easy and uncomplicated and a more widespread use would be a benefit for basic and applied research in the field.

Our bioinformatic analysis gives example that the rapidly increasing numbers of cyanobacterial genomes will help to identify potentially transformable species and offers a simple way to rank potential candidates. A recently published similar analysis (24) also comes to the conclusion that natural competence could be a common feature among cyanobacteria. However, not only the complete set of essential genes for natural competence but also their coordinated expression is important. Gene expression of NTFs is probably dependent on internal or environmental parameters. Studies on the regulation of natural competence are mostly limited to the gram positive genera *Bacillus* and *Streptococcus* and the gram negative *Vibrio cholerae* (48, 49).

All *Phormidium lacuna* filaments are highly motile on agar medium (35), and motility of cyanobacteria is thought to be dependent on type IV pili (50), the same structure that mediates NT. Motility indicates that *pil* genes are expressed, and a preselection of motile strains or conditions that increase motility could help in successful NT. The filamentous growth and motility on agar surfaces and different membranes is the reason why no colonies are obtained after transformation and the respective frequencies cannot be calculated as cfu/µg of plasmid DNA.

Genetic engineering is now possible for another cyanobacterium, which can be used in the future for basic research and biotechnological applications. The established protocol includes few washing steps, DNA addition and subcultivation, and clear results are obtained after about 4 weeks. Homogenization of filaments and subsequent washings could remove extracellular polymeric materials and nucleases (5) and could thereby promote NT. Members of the genus *Arthrospira* (or *Spirulina*), for which high nuclease activities were reported (40, 51), might be transformed by the same or a similar approach. Trials with other cyanobacteria that follow either the standard protocol or a simplified version of it are easy and can rapidly provide clear results.

The *Phormidium lacuna* transformants have an extraordinarily high resistance against Kn. We have in the meantime targeted other loci and found a strong antibiotic resistance for these transformants as well. We therefore would rule out a gene-positional effect on kanR expression. Although the origin of this strong resistance is unclear, it could accelerate the segregation into chromosomes and thereby speed up the entire transformation process.

We have shown that a newly isolated cyanobacterium *Phormidium lacuna* can be transformed by natural transformation and homologous recombination during which a kanR resistance cassette is integrated into the genome, that the transformants are extraordinarily resistant against Kn, that the cyanobacterium has multiple chromosome copies and that rapid segregation of the kanR resistance cassette into all chromosomes can be by achieved by the high resistance. We hope that our results stimulate trials on natural transformation of other cyanobacteria and thereby contribute to a broadening of research on more species, especially the filamentous ones.

## Acknowledgments

The work for supported by the PhD fellowship of the Nagelschneider Foundation to Fabian Nies. We thank Nadja Wunsch for technical assistance.

## Confilct of interest

The authors declare that there is no conflict of interest with other parties

